# Self-reported effects of classic psychedelics on stuttering

**DOI:** 10.1101/2023.04.18.537312

**Authors:** Noah D. Gold, Noam Goldway, Hope Gerlach-Houck, Eric S. Jackson

## Abstract

Stuttering is a neurodevelopmental communication disorder that can lead to significant social, occupational, and educational challenges. Traditional behavioral interventions for stuttering can be helpful, but effects are often limited. Classic psychedelics hold promise as a complement to traditional interventions, but their impact on stuttering is unknown. We conducted a qualitative content analysis to explore potential benefits and negative effects of psychedelics on stuttering using publicly available Reddit posts. A combined inductive-deductive approach was used whereby meaningful units were extracted and codes were initially assigned inductively. We then deductively applied an established framework to organize the effects (i.e., codes) into five subthemes (Behavioral, Emotional, Cognitive, Belief, and Social Connection), each of which was grouped under an organizing theme (positive, negative, neutral). Results indicated that the effects of psychedelics spanned all subthemes. Nearly 75% of participants reported overall positive effects. Nearly 60% of participants indicated positive behavioral change (e.g., reduced stuttering, increased speech control), 40% reported positive emotional benefit, 15% reported positive cognitive changes, 12% reported positive effects on beliefs, and 7% indicated positive social effects. Approximately 10% of participants reported negative behavioral effects (e.g., increased stuttering, reduced speech control). Psychedelics may help many stutterers improve communication, cultivate a healthier outlook, and promote psychological well-being. These preliminary results indicate that future clinical trials investigating psychedelic-assisted speech therapy for stuttering are warranted.

## 1. Introduction

Stuttering is a neurodevelopmental condition typically characterized by intermittent disruptions in speech production, including within-word repetitions and audible and inaudible sound prolongations, as well as accessory non-speech behaviors such as eye blinking, physical tension, and head jerking. Living with stuttering can have detrimental consequences on one’s quality of life as it is linked to adverse listener reactions, negative stereotypes, and significant educational and occupational disadvantages (Craig, Blumgart, & Tran, 2009; Gerlach, Totty, & Subramanian, 2018). The condition is common in early life, with about 5-8% of children exhibiting stuttering during the preschool years (Yairi & Ambrose, 2012). While most individuals recover spontaneously during early childhood, 20-25% of children continue to stutter into adulthood, making stuttering a common, lifelong condition that is present in about 1% of the adult population (Yairi & Ambrose, 2012).

While speech-related behaviors associated with stuttering (repetitions, prolongations) are most noticeable, affective and cognitive reactions such as anxiety, fear, and shame are primary contributors to the stuttering experience (Plexico et al., 2009b, 2009a; Tichenor & Yaruss, 2018, 2019b). This is due to negative listener reactions to stuttering (e.g., teasing, mocking, confusion), which contribute to social anxiety (Iverach et al., 2009, 2016) as stutterers learn to expect these reactions in their daily interactions. These expectations strengthen the anticipation of stuttering – the speaker’s ability to sense upcoming stuttering (Jackson et al., 2015) – which is linked to repetitive negative thinking or rumination (Tichenor & Yaruss, 2019a) (e.g., “I won’t be able to get this word out”; “People will think I’m stupid”). Such mental habits can lead to increased word or situational avoidance (Jackson et al., 2015) further amplifying one’s negative affective state. This complex interplay between social mistreatment and the behavioral, affective, and cognitive components of stuttering shapes the stutterer’s sense of self and identity (Daniels & Gabel, 2004; Gerlach et al., 2021) which is critical to address in therapy (Rodgers & Gerlach-Houck, 2022). Indeed, a significant component of clinical intervention for stuttering is based on counseling psychology and psychotherapeutic approaches, targeting affective-and cognitive-related aspects of stuttering and aiming to facilitate the change of maladaptive or rigid patterns of stutterers’ thoughts and behaviors. Stuttered speech is amenable to change in speech therapy, but such behavioral change is often not durable and old relics of tension, struggle, and avoidance while speaking often re-emerge (Craig & Calver, 1991; Craig & Hancock, 1995; Tichenor & Yaruss, 2020). In order for speech change to be durable, stutterers must also shift the way they identify with stuttering, which can involve exploring concepts such as openness and acceptance (Sisskin, 2023). Thus, new ways to achieve holistic improvement in stutterers’ well-being and quality of life, as well as increased ease of communication are critically needed.

In recent years, clinical trials have shown that psychedelic compounds have the potential to treat an array of psychiatric conditions that involve affective symptomatology (e.g., anxiety, depression) or rigid habitual behavioral patterns (e.g., addiction, obsessive-compulsive disorder) (Andersen et al., 2021; Leger & Unterwald, 2022; Nutt & Carhart-Harris, 2021). Classic psychedelics are a group of pharmacological agents defined by their high affinity for the 5HT_2A_ serotonin receptor and include drugs such as psilocybin, lysergic acid diethylamide (LSD), and dimethyl trypatime (DMT). Although the mechanisms of psychedelic-assisted treatments remain speculative (van Elk & Yaden, 2022), these drugs are thought to induce an acute period of psychological and cognitive flexibility (Davis et al., 2020; Doss et al., 2021; Pacheco et al., 2023) but see (Pokorny et al., 2020; Wießner et al., 2022), neural plasticity (de la Fuente Revenga et al., 2021; Ly et al., 2018; Raval et al., 2021; Shao et al., 2021), increased positive affect, and change in rigid patterns of maladaptive behavior (Goldberg et al., 2020). In their review, van Elk and Yaden (2022) proposed that the psychological effects associated with classic psychedelics comprise five categories: *behavior* (e.g., habit and behavior change); *altered and affective states* (e.g., ego dissolution, enhanced perceptions of emotions); *cognition* (e.g., cognitive flexibility, mindfulness); *beliefs* (e.g., metaphysical beliefs, suggestibility); and *social connection* (e.g., connectedness, empathy). Taken together, the evidence suggests that psychedelic compounds might complement existing behavioral therapies for stuttering with a specific emphasis on cognitive and affective symptomatology. However, with their potential benefits, psychedelics-assisted interventions carry potential psychiatric risks including acute mood instability and distortions in sensory perception, and long-lasting mania or psychosis (Bradberry et al., 2022; Johnson et al., 2008; Strassman, 1984). Such effects may be especially prevalent when dosing occurs in the absence of careful professional supervision and guidance (Johnson et al., 2008). Notably, numerous recent studies that examined the safety profile of psychedelics reported encouraging results (Bender & Hellerstein, 2022; e.g., Holze et al., 2022; Malcolm & Thomas, 2021). To date, there are no data regarding the safety profile of psychedelics in the context of stuttering.

Here, we investigated the types of experiences that individuals who stutter have when taking psychedelics in naturalistic settings. This was done in order to evaluate the potential therapeutic effects, and harmful effects, of classic psychedelics as a complement to traditional stuttering therapy. Psychedelics are being extensively used recreationally (Livne et al., 2022), and there is a lively online community that discusses various aspects of psychedelic use. In such online forums, individuals voluntarily share their experiences with psychedelic substances making them publicly available. We have utilized these resources to conduct qualitative and quantitative analyses of self-reported experiences of stutterers who have used psychedelics via posts on the online messaging forum *Reddit*.

## 2. Methods

We conducted a qualitative analysis by reviewing publicly available internet posts on the social news website, Reddit. Reddit is a well-known web-based platform where users post and have discussions about a wide variety of topics. Qualitative analysis was justified because we sought to develop a rich understanding of experiences with psychedelics and effects on stuttering among people who stutter, per their own report. We supplemented the qualitative analysis with descriptive statistics to assess how common various experiences with psychedelics are in this community. Such work may provide a foundation for future clinical trials on the efficacy of classic psychedelics on stuttering. Because all information was publicly available by an internet search, this study was exempt from Institutional Review Board approval. Participant demographics and related information were not obtained because Reddit is an anonymous forum.

### Author Positionality

Following a constructivist epistemology, our view is that truth and meaning in qualitative research cannot be completely independent of the authors’ experiences. Therefore, it is critical that all investigators reflect upon and report their positionality related to the research topic so readers can make their own inferences about potential influence. The first author is a research coordinator working on clinical trials utilizing psilocybin for the treatment of alcohol use disorder and other psychiatric indications. He has prior experience as a research assistant in a lab researching stuttering, and is a person who stutters. The second author is a post-doctoral fellow investigating the cognitive learning mechanisms that underlie psychiatric symptom expression. He is particularly interested in the cognitive mechanisms through which psychedelics extract their beneficial clinical effect. He is not a stutterer. The third author is a speech-language pathologist and researcher who studies how identity is influenced by living with stuttering using qualitative and quantitative methodologies. She is not a stutterer, and she has no prior personal or research experience with psychedelics. The last author is an Assistant Professor, speech-language pathologist with 15 years of experience and expertise in stuttering intervention, lab director, and person who stutters. His lab focuses on the variability of stuttering events, or the intermittency with which stuttering events occur, and the social-cognitive processes that drive this variability. He has been involved with stuttering and self-help communities since 2006, at which time he co-founded and co-hosted a podcast about stuttering which had the goal of reducing stigma associated with stuttering and normalizing stuttering.

### Data Acquisition

Self-report was used in the current study insofar as users self-identified as stutterers in their Reddit posts. Self-report is commonly used to identify stutterers in stuttering research (e.g., Gerlach et al., 2021). Combinations of the keywords, “stuttering,” “speech,” “psychedelics,” “psilocybin,” “mushrooms,” “LSD,” “DMT,” and “MDMA,” were used in the Reddit search. The keyword searches used the name of each psychedelic drug combined with either “stuttering” or “speech.” Although posts related to the effects of MDMA were not included in the analysis, MDMA was included as a search term because MDMA-related posts often included information about the use of classic psychedelics. While reviewing results within a Reddit discussion thread, both the original post as well as responses to the original post were reviewed for relevance to psychedelics and stuttering. Numerous replying posts (other Reddit users replying to the original post) included similar or shared experiences to that of the original post, so many of these were also included in the analysis. Posts were included for further review if they were 1) in English and comprehensible; and 2) included an account of direct first-hand experience on the effects of classic psychedelics on stuttering. Posts were copied in their entirety into a Microsoft Excel spreadsheet for analysis by the research team. All included data were posted on Reddit between 2012 and 2022.

### Data Analysis

Procedures from Braun & Clarke’s (2023; 2021) Reflexive Thematic Analysis were used to analyze and summarize meaning in the data. Reflexive Thematic Analysis relies on a constructivist epistemology and allows for inductive and deductive analysis approaches. The authors first familiarized themselves with the dataset. During this process, the first, second, and final authors extracted meaningful units from each post, or distinct utterances that were related to the question (i.e., the impact of classic psychedelics on stuttering). Text that was unrelated to the research question was excluded from the analysis. In addition, posts were removed if they did not describe experiences with the classic psychedelics (i.e., psilocybin, LSD, DMT), did not discuss personal, first-hand experiences, or included third-hand accounts or hearsay from another individual. Further, any duplicate posts (or those telling the same account, identified by the poster’s Reddit username) were removed. While MDMA was included in the original search terms because it sometimes yielded posts about classic psychedelics, meaningful units including only MDMA or other non-classic psychedelics were also excluded.

After all meaningful units had been identified and extracted, the first and last authors independently coded each meaningful unit with one or more codes using the constant comparative method (Glaser & Strauss, 2017). The constant comparative method involves iteratively comparing each meaningful unit to existing codes before assigning new ones, and updating code labels as nuance in the data becomes more apparent. The initial coding was inductive, in that it was not hypothesis driven, and we aimed to capture both semantic (i.e., explicit) and latent (i.e., implied) meaning (Braun & Clarke, 2021). Once both authors assigned one or more codes to all meaningful units they met to compare, discuss, and reach agreement on codes that best captured the content of each meaningful unit. As an additional effort to promote investigator triangulation, the second and third authors were invited to independently code the same meaningful units, and coding schemes were discussed once again until the authors agreed that all relevant meaning had been best captured with the codes.

Because we were interested in exploring *positive*, *negative*, and *neutral* effects of psychedelics on the experience of stuttering, we used these three categories as organizing themes as we continued to analyze the data. After the initial coding process was complete, we used a deductive approach to group codes into sub-themes, drawing from van Elk and Yaden’s (2022) framework on the five effects and therapeutic potential of psychedelics for theme content and organization (*Behavior, Altered and Affective States* [henceforth, *Emotion*], *Cognition, Beliefs,* and *Social Connection)*. For example, the codes “reduced stuttering” and, “improved speech” were grouped into the “Behavior” sub-theme, which fell under the broader theme of “Positive Effects,” whereas, “increased anxiety” and “increased confidence” were grouped into the “Emotions” sub-theme, which fell under the broader theme of “Negative Effects.” While certain code’s effects could be deemed positive vs. negative vs. neutral differently according to subjective preference of who is judging, valence was determined based on the third and fourth author’s clinical expertise in terms of what stutterers and the general public would deem as a positive or “beneficial” effect to someone who stutters. For example, reduced stuttering would be perceived by most individuals as a positive effect, increased anxiety as a negative effect, and no impact of psychedelics would be perceived as a neutral effect. Table 1 includes examples of meaningful units, their associated codes, and their broader thematic categories.

**Table 1:** Examples of meaningful units, and associated codes, subthemes, and themes.

| Meaningful Units | Code(s) | Subthemes | Themes |
| --- | --- | --- | --- |
| "The first time I ever microdosed (about 0.1g) my stutter was 99% non-existent." | Reduced Stuttering | Behavior | Positive |
| "I haven't noticed any difference in my stuttering before, after, or during shroom experiences." | No Effect on Stuttering | Behavior | Neutral |
| "Done a lot of psychedelics and they have not cured my stuttering but they're still fun and give you a little objectivity. One problem with a lot of stutterers is feeling anchored down in the moment by anxiety and a visceral sense of helplessness. Taking mushrooms and meditating helped a lot to let go of that anxiety which over the years has made stuttering a lot easier to deal with." | No Cure;<br>Increased Objectivity;<br>Reduced Anxiety;<br>Reduced Helplessness | Beliefs<br>Cognition<br>Emotion | Neutral<br>Positive |

To get a sense of the overall quality of individuals’ experiences with psychedelics, we included an additional level of analysis which coded the valence of each individual post holistically. For example, if most of the codes associated with a post were deemed positive, or the benefits of psychedelics were judged to be the focus of the post, despite their being some negative effect, the post was judged to be “positive.” Posts were judged to be “mixed” if there was approximately the same emphasis on positive and negative effects.

## 3. Results

Our final sample included 114 different Reddit posts from distinct Reddit users, totaling 709 sentences. Each of the 114 individuals contributed at least one meaningful unit, with 250 meaningful units in total. We assigned 77 unique codes across meaningful units, with 275 total codes assigned as meaningful units could have more than one code to describe effects of psychedelics. We developed three organizing themes, each with one or more subthemes, presented below.

### 3.1 Organizing Theme 1: Positive Effects

Two-hundred and sixteen codes were positively valenced (78.5% of our entire code sample), comprising the positive effects organizing theme. These meaningful units were further sub-grouped into five sub-themes described below. Table 2 includes all positive codes and their frequencies.

**Table 2.** Positively valenced codes and corresponding frequencies. % = percentage of all codes in the positive e thematic category; # = frequency; %* = percentage of all codes in the entire study; %** = percentage of all participants who reported this type of effect (# of individuals reporting this effect / 114 participants).

| Emotion | # | Cognition | # | Beliefs | # | Social | # | Behavior | # | Total |
| --- | --- | --- | --- | --- | --- | --- | --- | --- | --- | --- |
| Reduced Anxiety | 12 | Facilitated Introspection | 5 | Increased Acceptance | 7 | Reduced Social Anxiety | 2 | Reduced Stuttering | 54 |  |
| Improved Overall Wellbeing | 10 | Reduced Self-Consciousness | 4 | Increased Understanding of Oneself/One's Stuttering | 5 | Increased Ability to Communicate | 1 | Improved Speech | 5 |  |
| Improved Perspective | 9 | Increased Clarity | 3 | Hopeful | 1 | Bonded with Other Stutterers | 1 | Increased Control | 5 |  |
| Increased Confidence | 7 | Reduced Interfering Thoughts | 3 | Increased Appreciation of Life | 1 | Reduced Time Pressure | 1 | Increased Flow | 7 |  |
| Reduced Emotional Reaction | 5 | Increased Motivation | 2 | Lost Stutterer Identity | 1 |  |  | Reduced Effort | 5 |  |
| Reduced Ego | 4 | Potential to Disrupt Patterns of Behavior | 2 | Increased Authenticity | 1 |  |  | Reduced Avoidance | 3 |  |
| Reduced Concern about Stuttering | 3 | "Let Go" of Need for Control | 1 |  |  |  |  | Cured Stuttering | 2 |  |
| Increased Agency | 3 | Distracted Away from Stuttering | 1 |  |  |  |  | Changed How One Stutters | 1 |  |
| Psychedelics Helped | 3 | Facilitated Awareness of Behavior Patterns | 1 |  |  |  |  | Increased Body Relaxation | 1 |  |
| Reduced Stress | 3 | Increased Awareness of Speech | 1 |  |  |  |  |  |  |  |
| Reduced Anticipation | 2 | Increased Objectivity | 1 |  |  |  |  |  |  |  |
| Reduced Fear | 2 | Interrupted Unhelpful Feedback Loops | 1 |  |  |  |  |  |  |  |
| Increased Self-Esteem | 2 | Organization of Thought | 1 |  |  |  |  |  |  |  |
| Liberating | 2 | Prevented Snowball Effect | 1 |  |  |  |  |  |  |  |
| Afterglow Effect | 2 | Increased Mindfulness | 1 |  |  |  |  |  |  |  |
| Reduced Depression | 1 | Moved Past Mental Hang-Ups | 1 |  |  |  |  |  |  |  |
| Bypassed Shame and Fear | 1 | Facilitated Identification of Accessories | 1 |  |  |  |  |  |  |  |
| Improved Mood | 1 | Improved Language Ability | 1 |  |  |  |  |  |  |  |
| Personal Growth | 1 | Increased Clarity | 3 |  |  |  |  |  |  |  |
| Reduced Helplessness | 1 | Reduced Interfering Thoughts | 3 |  |  |  |  |  |  |  |
| Reduced Suffering | 1 |  |  |  |  |  |  |  |  |  |
| Reduced Trauma | 1 |  |  |  |  |  |  |  |  |  |
| Reduced Uncertainty | 1 |  |  |  |  |  |  |  |  |  |
| Opportunity to Accumulate Positive Experiences | 1 |  |  |  |  |  |  |  |  |  |
| Psychedelics Support Change, Which Helps Stuttering | 1 |  |  |  |  |  |  |  |  |  |
| Increased Naturalness | 1 |  |  |  |  |  |  |  |  |  |
| Increased Comfortability | 1 |  |  |  |  |  |  |  |  |  |
| **Total Count** | **81** |  | **31** |  | **16** |  | **5** |  | **83** | **216** |
| **%** | **37.50%** |  | **14.35%** |  | **7.41%** |  | **2.31%** |  | **38.43%** | **100.00%** |
| **%\* of all codes** | **29.45%** |  | **11.27%** |  | **5.81%** |  | **1.82%** |  | **30.18%** | **78.53%** |
| **%\*\* of participants** | **41.23%** |  | **18.42%** |  | **12.28%** |  | **4.39%** |  | **58.77%** | **N/A** |

#### 3.1.1 Subtheme 1a: Positive Behavioral Changes

Positively valenced effects on behavior were observed in 58.8% of individuals (67 out of 114 total individuals), who contributed a total of 83 codes (30.18% of the entire 275 code sample). The most commonly reported effect across all participants was a reduction in stuttering, reported by 54 individuals (47.4%). In addition, five individuals (4.4% of all individuals) commented on general improvements in their speech, and two individuals reported that psychedelics completely cured their stuttering. Individuals in our sample commonly cited that psychedelics caused reductions in their stuttering both in an acute manner while actively experiencing psychedelic effects as well as for varying amounts of time after. Twenty-seven individuals reported acute reductions only, 17 specified that they saw reductions persisting after their acute psychedelic effects wore off, and 10 reported reductions in stuttering without clearly specifying the length of this effect). Several individuals emphasized how they were “the most fluent that [they’ve] ever been” (P23 and P97). Another participant commented on being *completely* fluent, and attributed this to not being concerned about how they are perceived by others: Individuals also described that psychedelics helped to reduce their effort in speaking (4.4% of all individuals) and avoidance behaviors (2.6% of all individuals), and five individuals (4.4% of all individuals) reported increased control of their speech. In addition, seven individuals reported that psychedelics helped to improve their “flow” in speaking (6.1% of all individuals). For example, they cited that words “flowed out” (P9) easier, and they were able to “roll over [words] with much more ease” (P59). Other less commonly reported effects were increased body relaxation, and a change in “how” one stutters.

#### 3.1.2 Subtheme 1b: Positive Effects on Emotions

Positively valenced effects on emotions were observed in 41.2% of individuals (47 out of 114 total individuals), who contributed a total of 81 codes (29.5% of the entire 275 code sample). Twelve individuals described reduced anxiety, three described reductions in stress, and one described reductions in depression following psychedelic use. These reductions in psychological distress were both related directly to their stuttering, and also reported as more general improvements in their psychological state. For example, one individual (P93) reported a “lingering sense of calmness / lower anxiety after a trip, which does positively contribute to [their] fluency.” Another user (P34) reported that their “inherent anxiety around speaking was simply gone. [They] distinctly remember how the absence of anxiety felt.” Seven individuals cited increased feelings of confidence following their psychedelic experience, and five reported reduced negative emotional reactions. Ten users described that psychedelic use resulted in improvements in their general wellbeing. Another commonly reported effect was an improved sense of perspective (7.9% of all users). For example, one individual reported that psilocybin has been a:

*“key factor in what not only has increased my fluency as a speaker, but just overall who I am as a person the lens that I see life through… helped me in my journey allowing me to love and appreciate my struggles … including my stutter … I began to see that my value as a person has nothing to do with the way I talk, but who I really was deep down” (P16)*.

Four individuals described how psychedelics reduced their sense of ego. Three individuals reported reduced concern about their stuttering, two reported reduced feelings of anticipating that they might stutter, and two reported personal growth not necessarily related to speech. Additionally, two individuals emphasized how their psychedelic use was “liberating,” (P94, P97) three individuals described an increased sense of agency, and three individuals (P82, P90, P109) reported that psychedelics seem to “help” them more broadly (i.e., improve their wellbeing). Other less commonly reported positive effects on general wellbeing were, mood improvements, reduced fear, increased mindfulness, increased naturalness, increased comfortability, reduced emotional reaction, reduced helplessness, reduced suffering, reduced trauma, and reduced uncertainty.

#### 3.1.3 Subtheme 1c: Positive Changes in Cognition

Cognitive effects comprised changes in attention or cognitive control, as well as cognitive flexibility (i.e., the ability to adapt to changing demands or circumstance to reach a goal). Positively valenced effects on cognition were observed in 18.4% of individuals (21 out of 114 users), who contributed a total of 31 codes (11.27% of the entire 275 code sample). Five individuals (4.4% of all users) reported that their psychedelic experience helped to facilitate introspection, and three individuals (2.6% of all individuals) reported an increased sense of clarity (e.g., *“*the barrier between thought and speech was completely broken down, but yet (at least on the comedown) I was completely cogent”; P50). Four individuals (3.5% of all users) described how their psychedelic use decreased their self-consciousness. Three individuals (2.6% of all users) reported that psychedelics helped to reduce interfering, unwanted thoughts related to their speech and stuttering. For example, one individual stated that “psychedelics will make you not think about the small things like speech when there are way trippier things around you” (P24). Two individuals (2.6% of all individuals) reported that psychedelics have potential to help them change their patterns of behavior.

Moreover, at least one individual reported each of the following experiences: increased awareness of speech in a helpful way, improved language ability, increased awareness of behavioral patterns, increased ability to identify accessory behaviors to stuttering (i.e., eye-blinking, facial twitches, etc.), a helpful distractedness away from stuttering, increased objectivity, increased mindfulness, improved organization of thoughts, ability to “let go” the need for control, greater ability to move past mental hang ups, and that psychedelics help to prevent a “snowball effect” of bad experiences of stuttering.

#### 3.1.4 Subtheme 1d: Positive Effects on Beliefs

Changes in beliefs comprised changes to the beliefs system including worldview and self-identity. Positively valenced effects on beliefs were observed in 12.3% of individuals (14 out of 114 total users), who contributed a total of 16 codes (5.8% of the entire 275 code sample). The most common effect reported in this category was an increase in acceptance of one’s identity as a stutterer, which was reported by seven individuals (6.1% of all individuals). For example, one individual stated that “there is no cure, it is about accepting yourself as someone who stutters and then trying to control that as much as possible” (P47). Another individual described positive effects from their psychedelic experiences, and a corresponding broad hopefulness at the possibility of seeing positive effects from psychedelics in the future. Additionally, five individuals (4.4% of all users) reported that psychedelics helped to increase their understanding of themselves and their stuttering. Less commonly reported positive effects from psychedelics on beliefs were an increased appreciation of life, a loss of one’s stuttering identity, an increase in authenticity, and a sense of hopefulness.

#### 3.1.5 Subtheme 1e: Positive Social Effects

Positively valenced social effects were observed in 4.4% of individuals, who contributed a total of five codes (1.8% of the entire 275 code sample). Two individuals reported a decrease in social anxiety following their psychedelic experience. For example, one individual stated that they were “…more at ease in social settings and less self conscious. Stuttering less, and when [they] do [they’re] experiencing less negative thoughts around [themself]” (P102). Two participants reported that psychedelics were able to improve their ability to communicate with others, and help them make their points successfully and with less effort. Another individual stated that they have been able to “complete so many conversations and people have actually understood what [they’re] saying without repeating [themself]… it’s been such an experience” (P10). Less commonly reported positive social effects from psychedelic use were reduced feelings of time pressure in social interactions, and a beneficial opportunity to have a mutual psychedelic experience with similar others who stutter.

### 3.2 Organizing Theme 2: Negative Effects

Twenty-eight codes out of the sample of 275 were negatively valenced (10.2%). Table 3 summarizes these negatively valenced codes and their frequencies.

**Table 3.** Negatively valanced codes and corresponding frequencies. # = frequency; %* = percentage of all codes in the entire study; % = percentage of all codes in the negative thematic category; %** = percentage of all participants who reported this type of effect (# of individuals reporting this effect / 114 participants).

| Emotion | # | Cognition | # | Beliefs | # | Social | # | Behavior | # | Total |
| --- | --- | --- | --- | --- | --- | --- | --- | --- | --- | --- |
| Increased Concern about Stuttering | 1 | Reduced Language Ability | 3 | N/A |  | Difficulty Having a Conversation | 1 | Increased Stuttering | 13 |  |
| Negative Side Effects | 1 | Reduced Focus | 2 |  |  |  |  | Difficulty Speaking | 4 |  |
| Negative After-Effect | 1 | Increased Self-Consciousness | 1 |  |  |  |  | Increased Tension | 1 |  |
| **Total Count** | **3** |  | **6** |  |  |  | **1** |  | **18** | **28** |
| **%** | **10.71%** |  | **21.43%** |  |  |  | **3.57%** |  | **64.29%** | **100.00%** |
| **%\* of all codes** | **1.09%** |  | **2.18%** |  |  |  | **0.36%** |  | **6.55%** | **10.18%** |
| **%\*\* of participants** | **2.63%** |  | **5.26%** |  |  |  | **0.88%** |  | **14.91%** | **N/A** |

#### 3.2.1 Subtheme 2a: Negative Behavioral Change

Negatively valenced effects associated with behavior were observed in 14.91% of individuals (17 out of 114 total individuals), who contributed a total of 18 codes (6.55% of the entire 275 code sample). These thirteen individuals commented on psychedelics causing an increase in their stuttering (11.4% of all individuals), with four of those instances being only acute or transient increases (i.e., during the psychedelic experience), and one describing “worsened” stuttering until the “morning after” (P84). Another participant said that “psychedelics make it soooo hard for [them] to have a conversation” (P79). Four individuals described how psychedelics made it difficult for them to speak or say words. For example, one individual said that they “oftentimes couldn’t speak at all” while on a psychedelic (P15) One individual reported feeling an increase in physical tension (in their jaw) during their psychedelic experience (e.g.,“I usually do more LSD than shrooms but usually it increases my stuttering from the the jaw tension it causes” [P48]).

#### 3.2.2 Subtheme 2b: Negative Effects on Emotions

Negatively valenced effects on emotions were observed in 2.6% of individuals (three out of 114 total individuals), who contributed a total of four codes (1.1% of the entire 275 code sample). One individual commented on an increase in their concern and worry about their stuttering during their psychedelic experience. One individual (P42) described negative after effects from their psychedelic use during their “comedown” and their stuttering returning to baseline levels, while another (P61) described negative side-effects described broadly as “damage.”

#### 3.2.3 Subtheme 2c: Negative Changes in Cognition

Negatively valenced effects related to cognition were observed in 5.3% of individuals (6 out of 114 total individuals), who contributed a total of 6 codes (2.2% of the entire 275 code sample). Two individuals described experiencing reduced focus during their psychedelic experience, and two individuals reported impaired language ability. For example, P69 stated that they “…[they] can’t put a word or sentence together to save [their] life.” One individual described an increase in self-consciousness on psychedelics (i.e., “I was very self conscious about my speech”; P15).

#### 3.2.4 Subtheme 2d: Negative Effects on Beliefs

No individuals contributed negative codes to this category.

#### 3.2.5 Subtheme 2e: Negative Social Effects

Negatively valenced social effects were observed in one individual who contributed one code which described that psychedelics make it “soooo hard for [them] to have a conversation.”

### 3.3 Organizing Theme 3: Neutral Effects

There were 31 codes, 11.3% of our code sample, characterized as neutral. Table 4 summarizes these neutrally valence codes and their frequencies.

**Table 4.** Neutrally valenced codes and corresponding frequencies. # = frequency; % = percentage of all codes in the neutral thematic category; %* = percentage of all codes in the entire study; %** = percentage of all participants who reported this type of effect (# of individuals reporting this effect / 114 participants).

| Emotion | # | Cognition | # | Beliefs | # | Social | # | Behavior | # | Total |
| --- | --- | --- | --- | --- | --- | --- | --- | --- | --- | --- |
| N/A |  | No "aha"<br>Moment | 1 | No Cure | 14 | N/A |  | No Effect on<br>Stuttering | 16 |  |
| **Total Count** |  |  | **1** |  | **14** |  |  |  | **16** | **31** |
| **%** |  |  | **3.23%** |  | **45.16%** |  |  |  | **51.61%** | **100.00%** |
| **%\* of all**<br>**codes** |  |  | **0.36%** |  | **5.09%** |  |  |  | **5.82%** | **11.27%** |
| **%\*\* of**<br>**participants** |  |  | **0.88%** |  | **12.28%** |  |  |  | **14.04%** | **N/A** |

#### 3.3.1 Subtheme 3a: Neutral Behavioral Change

Neutrally valenced effects associated with behaviors were observed in 14.0% of individuals (16 out of 114 total individuals), who contributed a total of 16 codes (5.8% of the entire code sample). Sixteen individuals (or 14.0% of our entire sample) reported that their psychedelic use had no effect on their stuttering.

#### 3.3.2 Subtheme 3b: Neutral Effects on Emotions

No individuals contributed codes to this category.

#### 3.3.3 Subtheme 3c: Neutral Changes in Cognition

One individual described a neutral effect on cognition, i.e., that during their psychedelic experience, they “did not have a specific moment/realization” that they could pinpoint that gave them insight about their stuttering, although they highlighted how it was an “amazing experience” (#53).

#### 3.3.4 Subtheme 3d: Neutral Belief Changes

Neutrally valenced effects associated with beliefs were observed in 12.3% of individuals (14 out of 114 total users), who contributed a total of 14 codes (5.1% of the entire code sample). All individuals who reported neutrally valenced effects on beliefs (14 individuals) noted that their psychedelic experience did not “cure” their stuttering. For example, one individual stated:

*“I don’t think it’s a magic cure but it may help with side effects of stuttering like depression. You discover yourself and might even realize that it doesn’t matter that you stutter which might make speaking a little easier” (#87).*

Oftentimes, these codes were accompanied by similar praise of psychedelics helping their speech somewhat, or their outlook/psychological state related to their stuttering, but that their psychedelic experience was not a “cure all.”

#### 3.3.5 Subtheme 3e: Neutral Social Effects

No individuals contributed codes to this category.

### 3.4 Other Codes

In this section, we summarize other codes, or information that was potentially relevant (e.g., whether effects were acute or longer term; demographic information; an individual’s goals prior to using a psychedelic), but did not directly describe the effects of psychedelics on themselves and their experience with stuttering. There were a total of 45 other codes. Ten individuals described that they have “microdosed,” while six individuals emphasized how they experience different effects dependent upon the type of psychedelic that they take. Other less commonly reported effects not directly relevant to the research question include reduced effectiveness over repeated use, the individual having a high motivation to address their stuttering going into their psychedelic experience, the placebo effect, and uncertainty of correlation vs. causation of psychedelics generally.

### 3.5 Descriptive Analysis of Valence by Participant

Valences were assigned at an individual participant level for each overall post (positive, negative, neutral, or mixed). Of 114 participants, 71.9% indicated an overall positive experience; 11.4% indicated an overall negative experience; 12.3% indicated an overall neutral experience; and 4.4% indicated a mixed experience with roughly equal positive and negative components. Mixed experiences included approximately the same emphasis on positive and negative effects (e.g., “When I have taken mushrooms in the past, my stuttering is non existent…the feeling of coming down and the stuttering coming back is pretty terrible though”; P42).

## 4. Discussion

We conducted qualitative and quantitative content analysis of Reddit posts related to stutterers’ experiences with classic psychedelics. To our knowledge, this is the first study of any kind to systematically investigate possible effects, therapeutic or otherwise, of psychedelic compounds on stuttering. The majority of participants (approximately 72%) reported positive overall effects of psychedelics on their experience with stuttering, including changes across behavioral, emotional, cognitive, belief system, and social domains. This preliminary evidence suggests that testing the safety and feasibility of classic psychedelics in the context of stuttering could be an important step in the search for new treatments, or treatment adjuncts, for stuttering.

Approximately 60% of participants reported positive change in the behavioral domain, which included reduced stuttering, reduced effort, “improved” speech, and increased control. Reduced stuttering was reported by nearly 50% of the participants. All of these behavioral effects relate to increased functioning or facilitation of speech motor control, but it is unclear how psychedelics would facilitate these kinds of changes. One possible mechanism could be reduced involvement of cognitive control processes during speech production. Recent work points to overactivation in the cognitive control network of stutterers prior to stuttered vs. fluent speech (Jackson et al., 2022; Orpella et al., 2023). Moreover, internally-versus externally-directed focus is known to disrupt simple or automatized motor movements (Kal et al., 2013; McNevin et al., 2003; Wulf et al., 2001), and there is some evidence that this is the case in stuttering (Eichorn et al., 2016; Jackson et al., 2016). This is analogous to the *centipede effect* or *Humphey’s law*, whereby “consciously thinking about one’s performance of a task that involves automatic processing impairs one’s performance of it” (Colman, 2015, p. 3867). Psychedelics have been suggested to reduce the involvement of control processes across a variety of cognitive and perceptual domains (Barrett et al., 2018; Bouso et al., 2013; Quednow et al., 2012; Schmidt et al., 2018; Vollenweider et al., 2007). It could be that psychedelics dampen hyperactivation in these right hemisphere areas, which reduces stuttering or more likely makes it easier to move through stuttering events potentially by limiting interference of cognitive control on the speech-motor system (Alm, 2014; Jackson et al., 2022; Orpella et al., 2023).

More than half of the participants reported positive affective change, including reductions in anxiety and other emotional reactions to stuttering and listener reactions to stuttering. It is well known that anxiety negatively impacts quality of life in stutterers, who often report uncertainty and fear related to how they will be perceived by others (e.g., as unintelligent, not confident), and how stuttering will impact their success in professional and personal relationships (Boyle, 2015; Craig & Tran, 2014; Iverach & Rapee, 2013). Carhart-Harris and Friston (2019) propose that some psychopathologies such as depression and anxiety may develop because of maladaptive thoughts and behaviors, and associated negative self-perception, and/or fearful and pessimistic outlooks that become entrenched and resistant to change. Their theory can be applied to stuttering therapy. For example, cognitive restructuring is a critical component in the therapeutic process for stuttering. It involves developing alternative ways of thinking about one’s situation in order to facilitate a more agentic lifestyle (Manning, 2009). However, these changes can be difficult to achieve due to strong feelings of shame, embarrassment, anger, and fear. According to Carhart-Harris and Friston (2019), strongly held beliefs or entrenched ways of thinking become more flexible under psychedelics. This may allow for individuals to be open or receptive to new ideas, and not only to ideas that have been systematized through life experience. Stutterers often learn to expect negative reactions from listeners based on previous experiences, particularly during childhood, e.g., when they were teased or bullied for stuttering (Blood & Blood, 2016). Indeed, participants in the current study reported reduced anxiety, depression, stress, worry, and fear, improved overall wellbeing, increased confidence and better ability to cope with uncertainty, and an improved outlook on life under psychedelics. Thus, psychedelics may facilitate the therapeutic process for stutterers by allowing them to be more open and receptive to thinking about themselves, their stuttering, and their speaking partners in different, and more constructive ways.

Anxiety is also a primary contributor to the difficulty associated with implementing speaking strategies in “real world” settings. Stuttering is significantly reduced when social demands are low, such as speaking to one’s self or speaking to an infant (Alm, 2014; Jackson et al., 2021). Anecdotally, stutterers report being able to use speaking strategies during therapy sessions in which social demands are low, but often have significant difficulty when outside of therapy (e.g., at work, with potential romantic partners). Anxiety and repetitive negative thinking may reduce one’s ability to focus on physical speaking strategies in these more “real” situations. For example, many speaking strategies aim to facilitate the transition from the onset to the nucleus of a word (i.e., where stuttering occurs). This involves developing a deep awareness and focus on the placement and movement of the articulators. This awareness of the articulators allows for the modification of tensing or pressure, so that one can move through stuttering events, physically, more easily. Anxiety, repetitive negative thoughts, uncertainty, and fear, for example, make it difficult to focus on lessening tension during (or before) moments of stuttering–similar to how these cognitive states can make performing any motor task more difficult. Reducing anxiety, concern, and fear about stuttering, while increasing awareness, reducing interfering thoughts, and increasing mindfulness may make it easier to focus on modifying articulatory movements. Moreover, reduced ego, increased acceptance, and reduced self-consciousness may alleviate negative thoughts the stutterer has about how they are being perceived, which could also increase one’s ability to focus on speaking strategies.

Despite a large proportion of positive effects reported by the participants in our sample, a portion of participants reported negative effects. Thirteen participants or 11% of the sample reported increased stuttering or increased tension, and overall difficulty speaking. It is not currently understood why psychedelics would increase stuttering or overall speaking difficulty. Interestingly, 10 of the 13 participants reported acute effects, i.e., only during the psychedelic experience. The three participants who indicated carry-over difficulty reported effects that lasted for several hours after the experience or the following day. No participants reported longer-term negative effects.

Knowledge regarding the individual traits that may increase the probability of experiencing adverse effects in response to psychedelic substances is lacking. Gold-standard clinical practice considers a family history of schizophrenia or bipolar disorder as a risk factor for long-term adverse effects. However, the exclusion of such participants from most modern clinical trials has limited the quantification of such risk (Bradberry et al., 2022). In the current study, which was based on reports “from the wild,” participants did not report their background conditions systematically. It could be the case that some of the individuals contributing data for this study had a background of increased risk that might explain some proportion of the variance in individual responses. In addition, the environmental and social conditions in which the drug was taken, often referred to as set and setting, are known to heavily influence the qualities of the psychedelic experience (Carhart-Harris et al., 2018; Gukasyan & Nayak, 2022). Early theories suggest that psychedelics may amplify the contents of consciousness in a nonspecific manner (Leary, 1961). To this end, psychological preparation and coordination of expectations as well as the physical and social conditions wherein the drug is consumed can dramatically shape the effects of the acute experience. In the online sample discussed here, several participants pointed to the set or setting being the primary factor in determining whether there were negative effects. For example, one participant indicated that the frequency of stuttering was reduced with friends, but increased at a music festival at which he did not know anybody. This has important implications for potential psychedelic-assisted treatment such that the setting and the expertise of the therapist will play a critical role. That no participants indicated longer-term negative side effects further supports the notion that psychedelics may not be harmful to those who take them in a controlled setting and with a knowledgeable guide (i.e., therapist/facilitator).

### Directions for Future Research and Clinical Trials

We propose that the psychedelic-assisted psychotherapy model commonly used in modern clinical trials for an array of psychiatric conditions could be used for stuttering. In this model, participants undergo preparatory and integrative psychotherapy before and after receiving a dose of psilocybin (or other psychedelic compounds) in the presence of trained and licensed therapists (Guss et al., 2020). Participants in the current study took psychedelics in uncontrolled and non-therapeutic contexts.It is our hope that augmenting participants’ safety and providing the proper social and psychological support, in accordance with the gold-standard clinical practice, could maximize the potential of psychedelic-assisted therapy for stuttering. Such treatment, provided by a trained clinician will optimize experience integration and may capitalize on the induction of neurobehavioural flexibility to promote meaningful, potentially long-term clinical benefit. Recent clinical trials have begun using a psychedelic-assisted “group” therapy model, whereby individuals partake in combined or group therapy sessions (Trope et al., 2019). In a recent study investigating older long-term HIV/AIDS survivors, participants reported strong praise for the group therapy components of the intervention (Anderson et al., 2020). The benefits of group therapy for stuttering are well known (e.g., Byrd et al., 2018; Levy, 2018; Sisskin, 2018). Given the significant negative impact on communication, and that people who stutter often deal with shared and/or similar common experiences and difficulties, a group therapy model with psychedelic-assisted therapy may be particularly beneficial for this population.

## Conclusion

People who stutter are in need of more effective treatments to manage intermittent disruptions in speech communication and also provide relief from distress that accompanies their social experiences, such as anxiety and depression. To date, there are no FDA approved pharmacotherapies approved to treat stuttering. Considering the promising results from our qualitative analysis which demonstrates that the vast majority of individuals reported positive experiences of psychedelics on their stuttering and psychological well being, we believe that a future clinical trial investigating the efficacy of psychedelics as a complement to traditional therapies in this population is warranted.

## References

Alm, P. A. (2014). Stuttering in relation to anxiety, temperament, and personality: Review and analysis with focus on causality. Journal of Fluency Disorders, 40, 5–21.

Andersen, K. A., Carhart-Harris, R., Nutt, D. J., & Erritzoe, D. (2021). Therapeutic effects of classic serotonergic psychedelics: A systematic review of modern-era clinical studies. Acta Psychiatrica Scandinavica, 143(2), 101–118.

Anderson, B. T., Danforth, A., Daroff, R., Stauffer, C., Ekman, E., Agin-Liebes, G., Trope, A., Boden, M. T., Dilley, J., & Mitchell, J. (2020). Psilocybin-assisted group therapy for demoralized older long-term AIDS survivor men: An open-label safety and feasibility pilot study. EClinicalMedicine, 27, 100538.

Barrett, F. S., Carbonaro, T. M., Hurwitz, E., Johnson, M. W., & Griffiths, R. R. (2018). Double-blind comparison of the two hallucinogens psilocybin and dextromethorphan: Effects on cognition. Psychopharmacology, 235(10), 2915–2927.

Bender, D., & Hellerstein, D. J. (2022). Assessing the risk–benefit profile of classical psychedelics: A clinical review of second-wave psychedelic research. Psychopharmacology, 239(6), 1907–1932. https://doi.org/10.1007/s00213-021-06049-6

Blood, G. W., & Blood, I. M. (2016). Long-term consequences of childhood bullying in adults who stutter: Social anxiety, fear of negative evaluation, self-esteem, and satisfaction with life. Journal of Fluency Disorders, 50, 72–84.

Bouso, J. C., Fábregas, J. M., Antonijoan, R. M., Rodríguez-Fornells, A., & Riba, J. (2013). Acute effects of ayahuasca on neuropsychological performance: Differences in executive function between experienced and occasional users. Psychopharmacology, 230(3), 415–424.

Boyle, M. P. (2015). Relationships between psychosocial factors and quality of life for adults who stutter. American Journal of Speech-Language Pathology, 24(1), 1–12.

Bradberry, M. M., Gukasyan, N., & Raison, C. L. (2022). Toward Risk-Benefit Assessments in Psychedelic- and MDMA-Assisted Therapies. JAMA Psychiatry, 79(6), 525–527. https://doi.org/10.1001/jamapsychiatry.2022.0665

Braun, V., & Clarke, V. (2023). Toward good practice in thematic analysis: Avoiding common problems and be (com) ing a knowing researcher. International Journal of Transgender Health, 24(1), 1–6.

Byrd, C. T., Gkalitsiou, Z., Werle, D., & Coalson, G. A. (2018). Exploring the effectiveness of an intensive treatment program for school-age children who stutter, Camp Dream. Speak. Live.: A follow-up study. Seminars in Speech and Language, 39(05), 458–468.

Carhart-Harris, R. L., & Friston, K. (2019). REBUS and the anarchic brain: Toward a unified model of the brain action of psychedelics. Pharmacological Reviews, 71(3), 316–344.

Carhart-Harris, R. L., Roseman, L., Haijen, E., Erritzoe, D., Watts, R., Branchi, I., & Kaelen, M. (2018). Psychedelics and the essential importance of context. Journal of Psychopharmacology, 32(7), 725–731.

Clarke, V., & Braun, V. (2021). Thematic analysis: A practical guide. Thematic Analysis, 1–100.

Colman, A. M. (2015). A dictionary of psychology. Oxford quick reference.

Craig, A., & Calver, P. (1991). Following up on treated stutterers: Studies of perceptions of fluency and job status. Journal of Speech, Language, and Hearing Research, 34(2), 279–284.

Craig, A., & Hancock, K. (1995). Self-reported factors related to relapse following treatment for stuttering. Australian Journal of Human Communication Disorders, 23(1), 48–60.

Craig, A., & Tran, Y. (2014). Trait and social anxiety in adults with chronic stuttering: Conclusions following meta-analysis. Journal of Fluency Disorders, 40, 35–43. https://doi.org/10.1016/j.jfludis.2014.01.001

Daniels, D. E., & Gabel, R. M. (2004). The impact of stuttering on identity construction. Topics in Language Disorders, 24(3), 200–215.

Davis, A. K., Barrett, F. S., & Griffiths, R. R. (2020). Psychological flexibility mediates the relations between acute psychedelic effects and subjective decreases in depression and anxiety. Journal of Contextual Behavioral Science, 15, 39–45.

de la Fuente Revenga, M., Zhu, B., Guevara, C. A., Naler, L. B., Saunders, J. M., Zhou, Z., Toneatti, R., Sierra, S., Wolstenholme, J. T., & Beardsley, P. M. (2021). Prolonged epigenomic and synaptic plasticity alterations following single exposure to a psychedelic in mice. Cell Reports, 37(3), 109836.

Doss, M. K., Považan, M., Rosenberg, M. D., Sepeda, N. D., Davis, A. K., Finan, P. H., Smith, G. S., Pekar, J. J., Barker, P. B., & Griffiths, R. R. (2021). Psilocybin therapy increases cognitive and neural flexibility in patients with major depressive disorder. Translational Psychiatry, 11(1), 1–10.

Eichorn, N., Marton, K., Schwartz, R. G., Melara, R. D., & Pirutinsky, S. (2016). Does Working Memory Enhance or Interfere With Speech Fluency in Adults Who Do and Do Not Stutter? Evidence From a Dual-Task Paradigm. Journal of Speech, Language, and Hearing Research, 59(3), 415–429.

Gerlach, H., Chaudoir, S. R., & Zebrowski, P. M. (2021). Relationships between stigma-identity constructs and psychological health outcomes among adults who stutter. Journal of Fluency Disorders, 70, 105842.

Glaser, B. G., & Strauss, A. L. (2017). The discovery of grounded theory: Strategies for qualitative research. Routledge.

Goldberg, S. B., Shechet, B., Nicholas, C. R., Ng, C. W., Deole, G., Chen, Z., & Raison, C. L. (2020). Post-acute psychological effects of classical serotonergic psychedelics: A systematic review and meta-analysis. Psychological Medicine, 50(16), 2655–2666.

Gukasyan, N., & Nayak, S. M. (2022). Psychedelics, placebo effects, and set and setting: Insights from common factors theory of psychotherapy. Transcultural Psychiatry, 59(5), 652–664.

Guss, J., Krause, R., & Sloshower, J. (2020). The Yale manual for psilocybin-assisted therapy of depression (Using acceptance and commitment therapy as a therapeutic frame).

Holze, F., Caluori, T. V., Vizeli, P., & Liechti, M. E. (2022). Safety pharmacology of acute LSD administration in healthy subjects. Psychopharmacology, 239(6), 1893–1905.

Iverach, L., Jones, M., McLellan, L. F., Lyneham, H. J., Menzies, R. G., Onslow, M., & Rapee, R. M. (2016). Prevalence of anxiety disorders among children who stutter. Journal of Fluency Disorders, 49, 13–28.

Iverach, L., O’Brian, S., Jones, M., Block, S., Lincoln, M., Harrison, E., Hewat, S., Menzies, R. G., Packman, A., & Onslow, M. (2009). Prevalence of anxiety disorders among adults seeking speech therapy for stuttering. Journal of Anxiety Disorders, 23(7), 928–934.

Iverach, L., & Rapee, R. (2013). Social anxiety disorder and stuttering: Current status and future directions. Journal of Fluency Disorders, 40, 69–82.

Jackson, E. S., Dravida, S., Zhang, X., Noah, J. A., Gracco, V., & Hirsch, J. (2022). Activation in Right Dorsolateral Prefrontal Cortex Underlies Stuttering Anticipation. Neurobiology of Language, 1–95.

Jackson, E. S., Miller, L. R., Warner, H. J., & Yaruss, J. S. (2021). Adults who stutter do not stutter during private speech. Journal of Fluency Disorders, 70, 105878.

Jackson, E. S., Tiede, M., Beal, D., & Whalen, D. H. (2016). The Impact of Social–Cognitive Stress on Speech Variability, Determinism, and Stability in Adults Who Do and Do Not Stutter. Journal of Speech, Language, and Hearing Research, 59(6), 1295–1314.

Jackson, E. S., Yaruss, J. S., Quesal, R. W., Terranova, V., & Whalen, D. H. (2015). Responses of adults who stutter to the anticipation of stuttering. Journal of Fluency Disorders, 45, 38–51. https://doi.org/10.1016/j.jfludis.2015.05.002

Johnson, M. W., Richards, W. A., & Griffiths, R. R. (2008). Human hallucinogen research: Guidelines for safety. Journal of Psychopharmacology, 22(6), 603–620.

Kal, E. C., Van der Kamp, J., & Houdijk, H. (2013). External attentional focus enhances movement automatization: A comprehensive test of the constrained action hypothesis. Human Movement Science, 32(4), 527–539.

Leary, T. (1961). Drugs, set & suggestibility. Annual Meeting of the American Psychological Association, 6.

Leger, R. F., & Unterwald, E. M. (2022). Assessing the effects of methodological differences on outcomes in the use of psychedelics in the treatment of anxiety and depressive disorders: A systematic review and meta-analysis. Journal of Psychopharmacology, 36(1), 20–30.

Levy, C. (2018). Interiorised stuttering: A group therapy approach. In Stuttering Therapies (pp. 104–121). Routledge.

Livne, O., Shmulewitz, D., Walsh, C., & Hasin, D. S. (2022). Adolescent and adult time trends in US hallucinogen use, 2002–19: Any use, and use of ecstasy, LSD and PCP. Addiction, 117(12), 3099–3109. https://doi.org/10.1111/add.15987

Ly, C., Greb, A. C., Cameron, L. P., Wong, J. M., Barragan, E. V., Wilson, P. C., Burbach, K. F., Zarandi, S. S., Sood, A., & Paddy, M. R. (2018). Psychedelics promote structural and functional neural plasticity. Cell Reports, 23(11), 3170–3182.

Malcolm, B., & Thomas, K. (2021). Serotonin toxicity of serotonergic psychedelics. Psychopharmacology, 1–11.

Manning, W. H. (2009). Clinical decision making in fluency disorders. Cengage Learning.

McNevin, N. H., Shea, C. H., & Wulf, G. (2003). Increasing the distance of an external focus of attention enhances learning. Psychological Research, 67(1), 22–29.

Nutt, D., & Carhart-Harris, R. (2021). The current status of psychedelics in psychiatry. JAMA Psychiatry, 78(2), 121–122.

Orpella, J., Flick, G., Assaneo, F., Pylkkanen, L., Poeppel, D., & Jackson, E.S. (2023). Global Motor Inhibition Precedes Stuttering Events. BioRxiv.

Pacheco, A. T., Olson, R. J., Garza, G., & Moghaddam, B. (2023). *Acute psilocybin enhances cognitive flexibility in rats* (p. 2023.01.09.523291). bioRxiv. https://doi.org/10.1101/2023.01.09.523291

Plexico, L., Manning, W. H., & Levitt, H. (2009a). Coping responses by adults who stutter: Part I. Protecting the self and others. Journal of Fluency Disorders, 34(2), 87–107.

Plexico, L., Manning, W. H., & Levitt, H. (2009b). Coping responses by adults who stutter: Part II. Approaching the problem and achieving agency. Journal of Fluency Disorders, 34(2), 108–126.

Pokorny, T., Duerler, P., Seifritz, E., Vollenweider, F. X., & Preller, K. H. (2020). LSD acutely impairs working memory, executive functions, and cognitive flexibility, but not risk-based decision-making. Psychological Medicine, 50(13), 2255–2264.

Quednow, B. B., Kometer, M., Geyer, M. A., & Vollenweider, F. X. (2012). Psilocybin-induced deficits in automatic and controlled inhibition are attenuated by ketanserin in healthy human volunteers. Neuropsychopharmacology, 37(3), 630–640.

Raval, N. R., Johansen, A., Donovan, L. L., Ros, N. F., Ozenne, B., Hansen, H. D., & Knudsen, G. M. (2021). A single dose of psilocybin increases synaptic density and decreases 5-HT2A receptor density in the pig brain. International Journal of Molecular Sciences, 22(2), 835.

Schmidt, A., Müller, F., Lenz, C., Dolder, P. C., Schmid, Y., Zanchi, D., Lang, U. E., Liechti, M. E., & Borgwardt, S. (2018). Acute LSD effects on response inhibition neural networks. Psychological Medicine, 48(9), 1464–1473.

Shao, L.-X., Liao, C., Gregg, I., Davoudian, P. A., Savalia, N. K., Delagarza, K., & Kwan, A. C. (2021). Psilocybin induces rapid and persistent growth of dendritic spines in frontal cortex in vivo. Neuron, 109(16), 2535–2544. e4.

Sisskin, V. (2018). Avoidance reduction therapy for stuttering (ARTs). More than Fluency: The Social, Emotional, and Cognitive Dimensions of Stuttering, 157–186.

Strassman, R. J. (1984). Adverse reactions to psychedelic drugs. A review of the literature. J Nerv Ment Dis, 172(10), 577–595.

Tichenor, S. E., & Yaruss, J. S. (2018). A phenomenological analysis of the experience of stuttering. American Journal of Speech-Language Pathology, 27(3S), 1180–1194.

Tichenor, S. E., & Yaruss, J. S. (2019a). Repetitive Negative Thinking, Temperament, and Adverse Impact in Adults Who Stutter. American Journal of Speech-Language Pathology, 1–15.

Tichenor, S. E., & Yaruss, J. S. (2019b). Stuttering as defined by adults who stutter. Journal of Speech, Language, and Hearing Research, 62(12), 4356–4369. https://doi.org/10.1044/2019_JSLHR-19-00137

Tichenor, S. E., & Yaruss, J. S. (2020). Recovery and relapse: Perspectives from adults who stutter. Journal of Speech, Language, and Hearing Research, 63(7), 2162–2176.

Trope, A., Anderson, B. T., Hooker, A. R., Glick, G., Stauffer, C., & Woolley, J. D. (2019). Psychedelic-assisted group therapy: A systematic review. Journal of Psychoactive Drugs, 51(2), 174–188.

van Elk, M., & Yaden, D. B. (2022). Pharmacological, neural, and psychological mechanisms underlying psychedelics: A critical review. Neuroscience & Biobehavioral Reviews, 104793.

Vollenweider, F. X., Csomor, P. A., Knappe, B., Geyer, M. A., & Quednow, B. B. (2007). The effects of the preferential 5-HT2A agonist psilocybin on prepulse inhibition of startle in healthy human volunteers depend on interstimulus interval. Neuropsychopharmacology, 32(9), 1876–1887.

Wießner, I., Olivieri, R., Falchi, M., Palhano-Fontes, F., Maia, L. O., Feilding, A., Araujo, D. B., Ribeiro, S., & Tófoli, L. F. (2022). LSD, afterglow and hangover: Increased episodic memory and verbal fluency, decreased cognitive flexibility. European Neuropsychopharmacology, 58, 7–19.

Wulf, G., McNevin, N., & Shea, C. H. (2001). The automaticity of complex motor skill learning as a function of attentional focus. The Quarterly Journal of Experimental Psychology: Section A, 54(4), 1143–1154.

Yairi, E., & Ambrose, N. (2012). Epidemiology of Stuttering: 21st Century Advances. Journal of Fluency Disorders, 38(2), 66–87.

